# CRISPR/Cas9 knockout of *PtaAGL16* improves post-drought recovery and shoot vitality in hybrid poplar

**DOI:** 10.64898/2026.10.04.756531

**Authors:** Alexander Fendel, Sarah Wodtke, Matthias Fladung, Tobias Bruegmann

## Abstract

Drought stress severely limits tree growth and survival, a problem intensifying under climate change due to the rising frequency of extreme weather events. Identifying genetic variants that modulate drought stress responses and enhance resilience is therefore of great interest for developing climate-adapted trees. While the MADS-box transcription factor AGL16 has been implicated in drought responses and stomatal regulation in herbaceous species, the function of its homolog in woody perennials remains largely unexplored. Here, we investigate the role of PtaAGL16 in the poplar hybrid *Populus tremula* × *P. alba*. We generated CRISPR/Cas9 knockout lines of *PtaAGL16* in the poplar clone INRA 717-1B4. Plants were subjected to a severe drought stress regime followed by a re-watering recovery phase. Morphological and physiological traits, including growth, stomatal conductance, shoot vitality, and stomatal density, together with stress-associated metabolites and drought-responsive gene expression, were analyzed. All *agl16* knockout lines exhibited significantly reduced drought-induced damage compared to the wild-type. While the wild-type suffered extensive defoliation and persistent stem necrosis, the knockout lines retained higher shoot vitality and resumed growth after re-watering, including the formation of new leaves. Consequently, vitality in the knockout lines reached up to 86%, compared to only 34% in the wild-type. This improved resilience was accompanied by reduced persistent oxidative membrane damage, as indicated by lower MDA levels after recovery. These findings indicate that the disruption of *PtaAGL16* is associated with enhanced drought resilience in a woody perennial species and highlight its potential as a genetic target for improving climate adaptation in fast-growing forest trees.

## Introduction

Progressive climate change is affecting the vitality of forests and trees across Central Europe. Resulting long-lasting drought events in Europe have led to significantly reduced tree vitality, often characterised by early crown defoliation (Süßel and Brüggemann 2021; Rukh et al. 2023; Knutzen et al. 2025). In extremes, the loss of vitality continued significantly until dieback (Schuldt et al. 2020; Rukh et al. 2023). Even in the absence of mortality, drought still reduces tree growth and biomass production (Del Martinez Castillo et al. 2022; Machová et al. 2024) and renders trees more susceptible to biotic stress (Morcillo et al. 2019; Mbedzi and Naidoo 2025). The cumulative effect of these stressors is reduced forest productivity and increased tree mortality (Knutzen et al. 2025).

Trees, however, possess a suite of adaptive mechanisms to cope with drought stress through a combination of physiological and molecular responses, including stomatal closure, osmotic adjustment, and the accumulation of compounds such as ABA and proline (Sharma et al. 2019; Zulfiqar and Ashraf 2023). Stomatal closure, primarily mediated by ABA, is a key mechanism to reduce transpirational water loss (Cotelle and Leonhardt 2019). Here, ABA triggers a signaling cascade involving protein phosphatases and kinases, which ultimately leads to the phosphorylation and activation of anion channels such as Slow Anion Channel-Associated 1 (SLAC1) , thereby promoting stomatal closure (Becker et al. 2003; Geiger et al. 2009; Huang et al. 2019). The genes orchestrating these responses constitute candidate targets for enhancing drought tolerance in trees. The nature and extent of these responses vary depending on the species and the intensity of the stress (Mathes et al. 2024; Niemczyk et al. 2024). Not least because trees are almost undomesticated, there are also significant intraspecific variations in drought stress response, for example, between European provenances with different climatic conditions (Nguyen et al. 2017; Gazol et al. 2023).

*AGL16* encodes a MADS-box transcription factor (TF) of the MIKC type. As a class of regulatory elements, MADS-box TFs have a broad functional spectrum comprising, for example, 107 genes in *A*rabidopsis *thaliana* and 105 in *P*. *trichocarpa* (Ng and Yanofsky 2001). Individual MADS-box genes can even regulate seemingly different pathways such as reproductive and vegetative meristem development simultaneously (Melzer et al. 2008; Bruegmann and Fladung 2019). This also applies to AGL16, which binds to the promoters of over 2,000 genes, thereby influencing different pathways (Dong et al. 2023). Among the regulatory genes involved in the stomatal control and development, the *AGAMOUS-LIKE 16* (*AGL16*) has gained increasing attention, particularly after the accumulation of *AGL16* mRNA in the guard cells of stomata was discovered in *A. thaliana* (Alvarez-Buylla et al. 2000). Drought-stressed *agl16* mutants further show a markedly higher survival rate than wild-type (WT) plants, a phenotype that has been associated with altered expression of *STOMATAL DENSITY AND DISTRIBUTION 1* (*SDD1*) and consequently reduced stomatal density (Zhao et al. 2020).

In addition to developmental effects, AGL16 has been reported to regulate genes involved in ABA homeostasis (Zhao et al. 2020; Zhao et al. 2021). In *A. thaliana*, AGL16 represses the expression of *ABSCISIC ALDEHYDE OXIDASE 3* (*AAO3*), which catalyses the final step in ABA biosynthesis, and induces *ABA 8′-HYDROXYLASE* (*CYP707A3*), which catalyses the catabolic degradation of ABA (Nambara and Marion-Poll 2005; Umezawa et al. 2006; Zhao et al. 2020). Also, in cotton (*Gossypium hirsutum*), silencing of *GhAGL16* was reported to increase drought stress tolerance, accompanied by increased proline content and reduced malondialdehyde (MDA) content, whereas the transgenic overexpression in *A. thaliana* led to reduced survival rates after drought stress, reduced relative water content and increased stomatal density (Lei et al. 2024a).

While the function of AGL16 in drought responses has been characterized in herbaceous species and cotton, the role of its homolog in long-lived woody tree species remains unknown. In the present study, we therefore disrupted *AGL16* in the poplar hybrid *Populus tremula* × *Populus alba* (often referred to as *Populus* × *canescens*), which serves as a model plant species for forest biotechnology (Leple et al. 1992; Bruegmann et al. 2019a). We generated CRISPR/Cas9 loss-of-function mutations in the *PtaAGL16_2* paralog and subjected three independently edited lines to a controlled drought stress regime followed by a recovery phase. The knockout (KO) of *PtaAGL16_2* was associated with enhanced drought resilience, as indicated by higher shoot vitality, preserved growth capacity, and lower persistent oxidative damage markers after re-watering.

## Materials and Methods

### Plant material and *in vitro* growth conditions

The poplar hybrid clone INRA 717-1B4 (*P. tremula* × *P. alba*; (Leple et al. 1992)) served as experimental material. WT and transgenic lines were maintained and propagated *in vitro* on solid McCown Woody Plant Medium (WPM; Duchefa, Haarlem, The Netherlands) containing vitamins, 2% (w/v) sucrose, and 0.6% (w/v) agar, adjusted to pH 5.8. Cultures were grown in Magenta vessels under controlled environmental conditions (23 °C, approx. 50% relative humidity) and exposed to continuous light (24 h) with a photosynthetic photon flux density (PPFD) of approx. 16 µmol m⁻² s⁻¹.

### Identification of *PtaAGL16,* and generation of CRISPR/Cas9 KO lines

To identify the *PtaAGL16* gene in *P*. *tremula* × *P*. *alba*, the *A. thaliana At*AGL16 protein sequence (AT3G57230) was used as a query in a BLAST search against the *P*. *tremula* × *P*. *alba* HAP1 v5.1 and HAP2 v5.1 genomes accessed via the Phytozome database (Zhou et al. 2023). The identified paralogous genes were located on chromosome 2 (PtXaTreH.02G093900, PtXaAlbH.02G094700) and chromosome 5 (PtXaTreH.05G115600, PtXaAlbH.05G117500). Spacers for CRISPR/Cas9-targeting gRNAs were manually selected based on the genomic sequences of the four identified *PtaAGL16* candidate genes. To ensure the functional KO, target sites were located within or upstream of the MADS domain, identified using the SMART web resource (Letunic et al. 2021). One spacer was designed to target both genes on chromosome 2, and a second spacer to target both genes on chromosome 5. The secondary structure of the full-length gRNA was predicted using the RNAfold WebServer (Gruber et al. 2008) and evaluated according to Bruegmann et al. (2019a). The CRISPR/Cas9 vector for *PtaAGL16* KO was obtained from DNA Cloning Service (Hamburg, Germany; Fig. S1). The T-DNA region contained the 35S::*hptII* selection marker gene, the 35S::*SpCas9* cassette, and two gRNAs driven by the *At*U6 promoter. The recombinant plasmid was introduced into *Agrobacterium tumefaciens* strain GV3101-pMP90RK and used to transform the WT poplar via a modified leaf disc method (Bruegmann et al. 2019b). Transformed poplars were regenerated on WPM supplemented with cefotaxime (500 mg L⁻¹) to eliminate residual *A. tumefaciens*, hygromycin (20 mg L⁻¹) for selection, as well as Pluronic F-68 (100 mg L⁻¹) and thidiazuron (0.0022 mg L⁻¹) to promote regeneration.

### Confirmation of transgenic status and genetic analyses in poplar lines

Genomic DNA was isolated from selected plant regenerates according to Bruegmann et al. (2022). Putative transgenic lines were confirmed by standard PCR experiments targeting the *hptII* and *SpCas9* genes. The CRISPR target regions were amplified and Sanger sequenced (StarSeq, Mainz, Germany) to detect genetic modifications. A Southern blot analysis was performed to determine the number of integrated T-DNA sequences in the putative KO lines (Fladung and Ahuja 1995). Here, at least 20 µg of genomic DNA was digested with *Dra*I. Further, possible off-target editing sites were identified with Cas-OFFinder (Bae et al. 2014). Off-targets with up to two mismatches between spacers and genomic sequences, which were located in gene-coding regions, were checked by PCR amplification and sequencing. T-DNA integration loci were identified using Thermal Asymmetric Interlaced PCR (TAIL-PCR) as described by Liu et al. (1995) and Fladung and Polak (2012). The resulting integration locus sequences were compared to the two *P. tremula* × *P. alba* haplotypes (HAP1 and HAP2) via BLAST analysis using the Phytozome database. All PCR primers are listed in Table S1. Based on these analyses, three transgenic lines displaying distinct genetic modification patterns were selected for further experiments: *agl16-3*, *agl16-7*, and *agl16-10*.

### Drought stress experimental setup

Rooted head cuttings were transferred from *in vitro* culture to propagation trays, filled with sieved tree nursery soil. Plants were acclimatized in a sealed container within a climate chamber (Weiss Technik, Lindenstruth, Germany) under a 16/8 h light/dark cycle (16 µmol m⁻² s⁻¹ PPFD, 22/17 °C, 70% RH) by gradually opening the lid over 10 days. Subsequently, plantlets were repotted into 1 L square plastic cultivation pots (top 11 × 11 cm, height 12 cm), filled with coarsely sieved tree nursery soil for optimal within-pot water distribution and the reduction of confounding substrate water-retention effects. Within the last three weeks of cultivation, all plants were fertilized once per week using 0.2% liquid fertilizer (WUXAL all-purpose fertilizer; Hauert MANNA Düngerwerke, Nuremberg, Germany). After 84 days of cultivation, the plants reached a height range of approx. 25 to 35 cm. Following, a three-week climate chamber experiment was conducted from November 26 to December 17, 2024. To minimize variability, we selected only plants with uniform growth and similar morphological characteristics for the drought experiment. In total, 40 plants (10 plants per line) were arranged randomized under the described environmental conditions, except PPFD increased to approx. 127 µmol m⁻² s⁻¹. For each line, five individuals were subjected to drought stress by withholding water for 7 days, followed by 14 days of re-watering to allow recovery. The remaining five plants per line served as well-watered controls, receiving 100 mL of water per day. Plants were phenotyped weekly and samples were collected at three time points (sampling days, SDs): start of the experiment (SD I), after 7 days of drought stress (SD II), and following 14 days of re-watering (SD III).

### Measurement of morphological, physiological, and molecular indicators

#### Morphological phenotyping

Plant height was measured from the stem base to the shoot apex. Stem diameter was recorded at 10 cm above the base. At each SD, fully expanded, vital leaves were counted for each plant. A leaf was defined as fully expanded when it reached a diameter of ≥ 2.5 cm (typically the 2^nd^ or 3^rd^ leaf from the apex) and considered vital if ≥ 90% of the leaf area consisted of chlorophyll-containing cells. To monitor new leaf formation, shoot tips located above the first fully expanded leaf were labeled following each sampling event. Shoot vitality was assessed by measuring the length of the visually green, chlorophyll-containing stem section, and the proportion of this length relative to total plant height was calculated and expressed as a percentage.

#### Stomatal density

Trichome removal and stomatal counting were performed as described by Wodtke et al. (2025). For each line, the third fully expanded leaf was collected from each individual plant at SD II for both treatments. Stomatal density (number per mm²) was determined by visualizing stomata with a light microscope (Axioscope 5; ZEISS, Oberkochen, Germany) at 20× magnification and counting using ImageJ 1.54p software (Schneider et al. 2012). Three positions per leaf were counted and averaged to obtain a single stomatal density value per leaf.

#### Stomatal conductance

Stomatal conductance was measured using an SC-1 leaf porometer (METER Group, Pullman, WA, USA). Measurements were taken on the 3^rd^ fully expanded leaf of each plant.

#### Molecular physiological phenotyping

Proline content was determined in the fourth fully expanded leaf, following the protocol of Bates et al. (1973) with modifications for a microplate-based assay (Abrahám et al. 2010). Leaf samples were collected around 10:00 a.m., immediately flash-frozen in liquid nitrogen, and stored at -80 °C until analysis. For the assay, 50 mg of leaf tissue was ground under liquid nitrogen cooling using a Bead Ruptor Elite homogenizer (Omni International, Kennesaw, GA, USA). To remove interfering proteins, each sample was combined with 1 mL of 5-sulfosalicylic acid and centrifuged at 12,000 × g for 10 min. An aliquot of 200 µL of the resulting supernatant was mixed with 200 µL of an acid-ninhydrin reagent (2.53% ninhydrin dissolved in a 3:2 (v/v) mixture of glacial acetic acid and 6 M phosphoric acid) and incubated at 96 °C for one hour. The reaction was terminated by placing the samples on ice, after which 1 mL of toluene was added to each tube to extract the proline-ninhydrin chromophore. Proline quantification was carried out by measuring the absorbance of the toluene phase at 520 nm in a 96-well plate using a microplate reader (SPECTROstar Nano; BMG LABTECH, Ortenberg, Germany). Each sample was analyzed in triplicate. A calibration curve was prepared with L-proline standards ranging from 0 to 50 µg mL⁻¹ (R² ≥ 0.985). The proline content, expressed in µmol per gram fresh weight (µmol g^−1^ FW), was calculated using the formula: (*C* × V) / (FW × 115.13), where *C* is the proline concentration derived from the standard curve, V is the total extraction volume (1 mL), FW is the sample fresh weight (0.05 g), and 115.13 g mol⁻¹ is the molecular weight of L-proline.

Lipid peroxidation was assessed by measuring MDA content, following the method described by Hodges et al. (1999). A 100 mg sample of the fourth fully expanded leaf was homogenized using the same procedure as for proline analysis, then mixed with 1.5 mL of 0.1% (w/v) trichloroacetic acid to precipitate proteins, and centrifuged at 14,000 × g and 4 °C for 15 min. Subsequently, 500 µL of the supernatant was combined in a 1:1 ratio with reaction solution 1 (RS1; containing 0.01% (w/v) 2,6-di-tert-butyl-4-methylphenol and 20% (w/v) TCA), while a parallel 500 µL aliquot was mixed with reaction solution 2 (RS2; RS1 supplemented with 0.65% (w/v) 2-thiobarbituric acid). Both mixtures were incubated at 96 °C for 30 min. The reactions were halted by placing the samples on ice, followed by centrifugation at 8,000 × g and 4 °C for 10 min. Absorbance was then measured at 440, 532, and 600 nm using the microplate reader, with 100 µL per well in three technical replicates for all samples and blanks (0.1% TCA combined with either RS1 or RS2). MDA concentrations were calculated using the equations provided by Hodges et al. (1999) and Landi (2017), and expressed as nmol per gram fresh weight (nmol g⁻¹ FW).

### Gene expression analyses

Real-time quantitative PCR (RT-qPCR) was used to analyze the expression levels of *PtaAGL16_2*, *PtaAGL16_5*, and selected antioxidant-, ABA-, and stomatal regulation-related genes (*PtaCAT2*, *PtaNCED3*, *PtaSDD1*, and *PtaSLAC1*; gene identifier in Table S1) in WT and the three KO lines. Total RNA was isolated from leaf tissue harvested on SD II between 11:00 a.m. and 12:00 p.m., using four biological replicates per line. RNA extraction was performed using the innuPREP Plant RNA Kit (IST Innuscreen, Berlin, Germany) according to Bruegmann et al. (2024). cDNA synthesis and RT-qPCR were carried out with the GoTaq 2-Step RT-qPCR System (Promega Corp., Madison, WI, USA) following the manufacturer’s instructions. Reactions were run on a C1000 Touch Thermal Cycler (Bio-Rad Laboratories Inc., Hercules, CA, USA) with the following profile: initial denaturation at 95 °C for 2 min, followed by 40 cycles of 95 °C for 15 s and target-specific annealing temperatures (*T*_A_, averaging 59 °C) for 1 min. Specific amplification was verified by melting curve analysis (95 °C for 15 s, followed by a 1 min step at the respective *T*_A_)). Expression levels were normalized to the poplar reference gene *UBIQUITIN* (*PtaUBQ*; Regier and Frey (2010)). ΔΔCt values were calculated using WT control as the single calibrator across all samples. Outliers were identified within each Line × Treatment group based on a median-distance criterion, whereby a sample was classified as an outlier if its ΔΔCt value differed by >1.5 Ct from the group median while the remaining biological replicates were within a ΔΔCt range of ≤1.0 Ct. Statistical analyses were performed on ΔΔCt values using a two-factor linear model (ΔΔCt ∼ Line × Treatment). Contrasts were computed from estimated marginal means (emmeans), including line differences within treatment, stress effects within line, and genotype–specific stress responses. *P*-values were FDR-adjusted (False Discovery Rate, Benjamini-Hochberg). Effects are reported as fold–change ratios after back–transformation to the2^−ΔΔCt^ scale (Livak and Schmittgen 2001).

### Statistical analyses of the drought stress experimental data

All analyses were performed in R version 4.5.2 and RStudio version 2026.1.0.392. Plants from different lines were measured under control and drought conditions across multiple SDs. Repeated measurements were accounted for by including plant ID as a random effect where applicable. For continuous traits measured over multiple SDs, linear mixed-effects models were fitted using the formula: *response ∼ Line × Treatment × SD + (1|ID)* (package *lmerTest*). Model assumptions were evaluated using residual diagnostics, and log(x + 1) transformations were applied where appropriate. Count traits were analyzed using generalized mixed-effects models with Poisson or negative binomial distributions, and when excess zeros affected model performance, hurdle models were applied. Proportions were analyzed using beta regression (package *glmmTMB*). End-point continuous traits were analyzed using linear models (*Line × Treatment*). Estimated marginal means and planned contrasts (line comparisons within *Treatment × SD*) were computed using the *emmeans* package with Holm correction applied for multiple testing. Diagnostics for generalized models were based on DHARMa residual simulations.

## Results

Two gRNA spacers were designed targeting the two *PtaAGL16* homologs, designated as *PtaAGL16_2* and *PtaAGL16_5,* located on chromosomes 2 and 5, respectively (Fig. 1a). The database gene annotation revealed that the *PtaAGL16_2* gene consists of seven exons, spanning a genomic region of 19,258 bp in the *P. tremula* allele and 20,364 bp in the *P. alba* allele with a notably large intron separating exons 1 and 2. The *PtaAGL16_5* spans a genomic region of 12,923 bp in the *P. tremula* allele and 13,103 bp in the *P. alba* allele with 8 exons. *In silico* analysis identified two protein domains (MADS and K-box) essential for the functionality of *PtaAGL16* as TF. To maximize the likelihood of loss-of-function mutations, the spacer sequences were designed to target the 5’ end of the first exon, targeting the MADS motif, which is critical for the DNA-binding capacity and functionality of the translated protein.

**Fig. 1.**
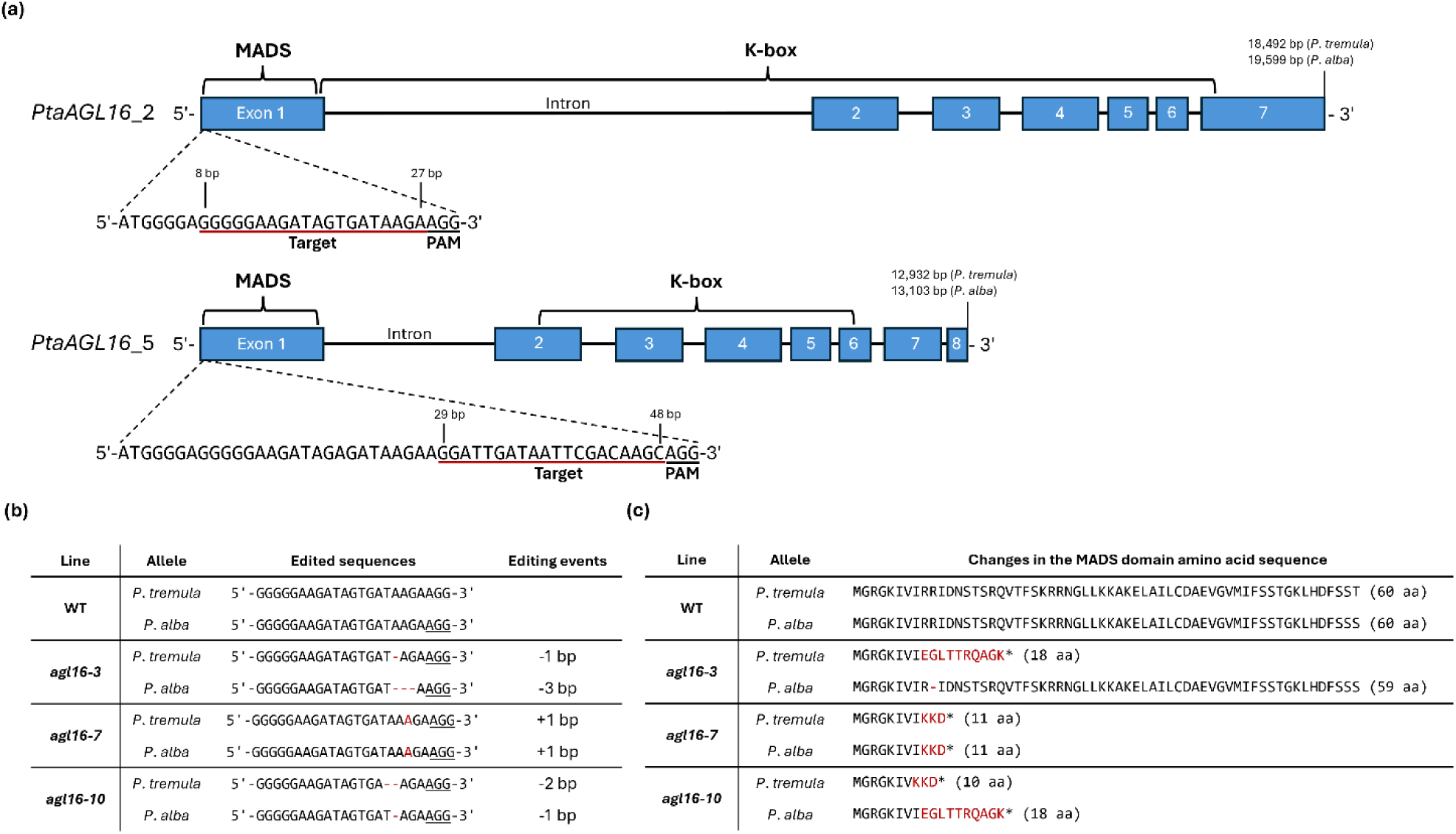
Design and results of CRISPR/Cas9 KO approach. (a) Genetic structure of the target genes *PtaAGL16_2* and *PtaAGL16_5* showing the number of exons (blue rectangles) and the location of the MADS and K-box protein motifs. The CRISPR/Cas9 targets and the corresponding PAM sequences are indicated in exon 1. (b) Editing events observed in the PtaAGL16_2 target, KO lines *agl16*-*3*, *agl16*-*7*, and *agl16*-*10* for each allele. (c) *In silico* predicted impact of indels on the *PtaAGL16_2* MADS domain translation. * – premature stop codon

Fifty-one regenerants were obtained following poplar transformation, of which 32 survived antibiotic selection. Ten of these were selected based on transformation-positive screening results and subjected to Sanger sequencing to obtain their gene editing pattern. Because the gRNA targeting the allele on chromosome 5 failed to induce editing, these loci remained unmodified; therefore, we focus exclusively on the *PtaAGL16_2* locus on chromosome 2 hereafter. Three lines (*agl16-3*, *agl16-7*, and *agl16-10*) exhibiting distinct modifications at the target site were chosen for further molecular analyses and the subsequent stress experiment. The genome editing outcomes varied among alleles: line *agl16-3* displayed a biallelic editing event, characterized by a 1 bp and 3 bp deletion in the *P. tremula* and *P. alba* alleles, respectively (Fig. 1b). Line *agl16-7* exhibited a homozygous editing event with a 1 bp insertion in both alleles, while *agl16-10* showed a biallelic event with a 2 bp deletion in the *P. tremula* allele and a 1 bp deletion in the *P. alba* allele. Southern blot analyses revealed a single T-DNA integration in all three lines, and subsequent TAIL-PCR followed by Sanger sequencing identified exclusively intergenic T-DNA insertion sites (Fig. S2; Table S2).

*In silico* modeling of the edited alleles predicted that most modifications result in premature translation termination due to frameshifts and the introduction of premature stop codons (Fig. 1c). Specifically, the homozygous editing in *agl16-7* led to stop codons after 11 amino acids (aa). Similarly, *agl16-10* contained premature stop codons after 10 aa in the *P. tremula* allele and 18 aa in the *P. alba* allele. A 3 bp deletion in the *P. alba* allele of *agl16-3* resulted in an *in-frame* deletion, removing only a single arginine residue from the MADS domain. In contrast, the *P. tremula* allele of this line harbored a premature stop codon after 18 aa. Collectively, these alterations are predicted to significantly compromise the functionality of the *PtaAGL16_2* TF. The complete coding sequences (CDS) of both WT alleles of *PtaAGL16_2* are shown in Fig. S3.

The drought stress experiment was conducted for three weeks in a climate chamber under constant environmental conditions and included the WT alongside the three KO lines. For each line, five individuals were assigned to either the control or the drought treatment. Under control conditions, morphological measurements indicated steady growth across all lines. Over the course of the experiment, shoot height increased by an average of 15.9 cm in the WT, and 16.3 cm, 14.3 cm, and 14.7 cm in *agl16-3*, *agl16-7*, and *agl16-10*, respectively (Fig. 2a). Similarly, stem diameter increased by approximately 1.00 cm in the WT, compared to 0.92 cm, 0.80 cm, and 0.75 cm in the respective mutant lines (Fig. 2b). Notably, significant differences in growth were observed among the lines. Line *agl16-7* exhibited significantly reduced shoot height compared to the WT at SD II (*p* < 0.01), a difference that persisted during the recovery phase at SD III. Similarly, *agl16-10* showed significantly lower shoot height than the WT (*p* < 0.05). Regarding stem diameter, only *agl16-7* displayed a narrower diameter compared to the WT at SD III (*p* < 0.05). In contrast, distinct differences were evident in the stress-treated plants. Between the start of the experiment and the severe stress on SD II, growth was generally inhibited in all lines. Here, WT (*p* < 0.001), *agl16-7* (*p* < 0.05), and *agl16-10* (*p* < 0.05) differed significantly from their respective controls. In the WT, growth arrest persisted during the recovery phase. In contrast, all three KO lines exhibited significant increases in shoot height, with an average gain of 4.6 cm over the 14 days of recovery at SD III (*agl16-3*: *p* < 0.001; *agl16-7*: *p* < 0.001; *agl16-10*: *p* < 0.01). The growth of *agl16-3* exceeded that of the WT (*p* < 0.01) and *agl16-7* (*p* < 0.05). Compared to their respective controls, all lines showed a smaller shoot height on SD III (*p* < 0.001). Stem diameter exhibited a similar pattern of growth arrest; however, recovery occurred more slowly than for shoot height and did not result in significant differences on SD III.

**Fig. 2.**
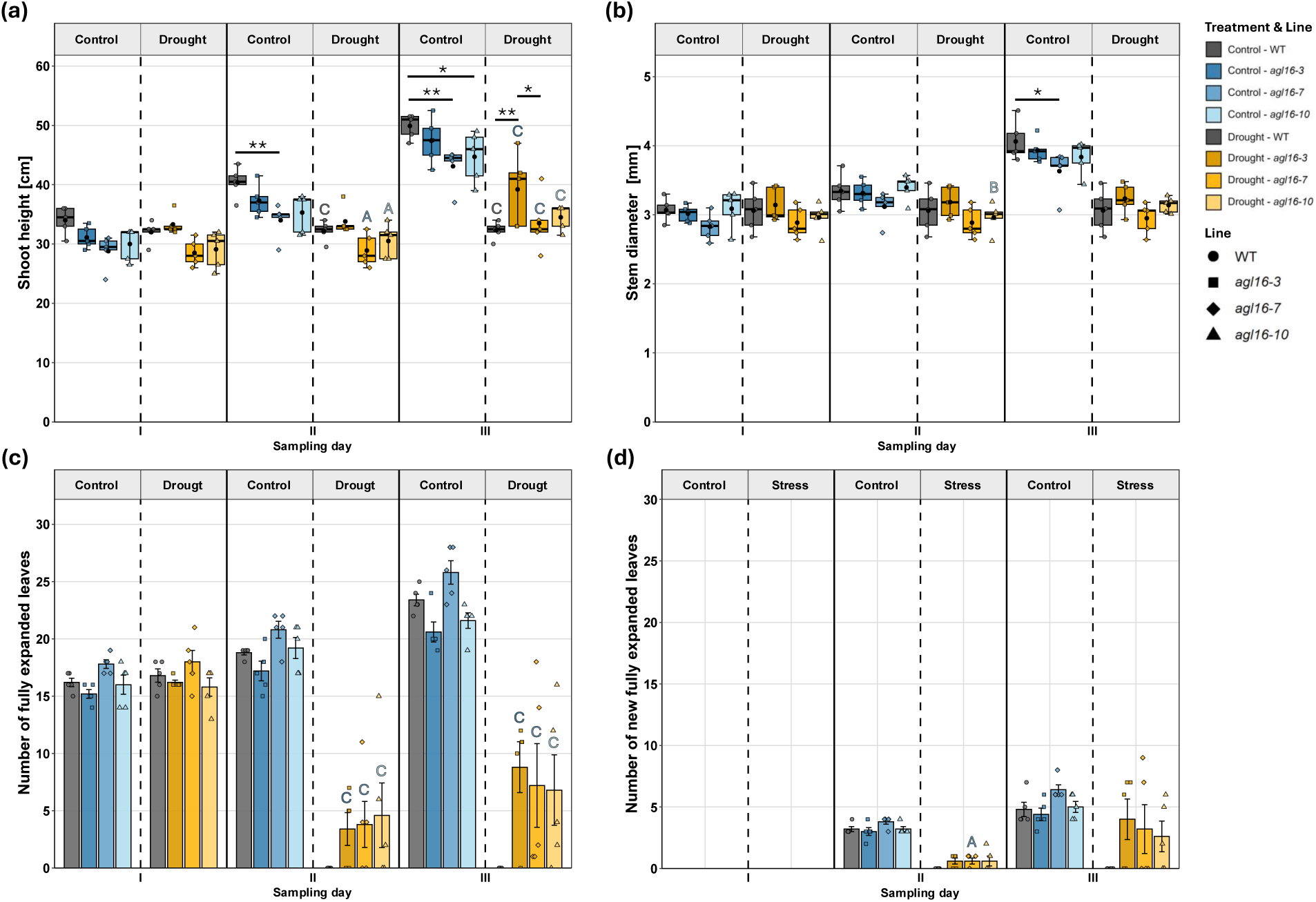
Morphological responses of WT and *PtaAGL16*_2 KO lines (*agl16*-*3*, *agl16*-*7*, *agl16*-*10*) to control and drought stress conditions. Measurements of (a) shoot height, (b) stem diameter, (c) number of fully expanded leaves, and (d) number of new fully expanded leaves across SDs (I, II, III) under control conditions and drought stress treatment. Data points visualize individual plants (n = 5 biological replicates per line × treatment × SD). For continuous traits (A, B), boxplots indicate the median and interquartile range (whiskers: 1.5× IQR) and black dots indicate means. For count traits (C, D), bars indicate means ± standard error. Asterisks and connecting lines denote significant pairwise differences among lines within each treatment × SD combination based on mixed-effects models followed by Holm-adjusted post hoc contrasts (* *p* < 0.05, ** *p* < 0.01, *** *p* < 0.001). Different capital letters above the stress groups indicate significant treatment contrasts within each line (*p* < 0.05, *p* < 0.01, *p* < 0.001, indicated by A, B, C, respectively). Letters are colored according to the respective control group of each line

To further assess growth performance, both the absolute leaf number and the number of new fully expanded leaves were recorded at each SD (Fig. 2c, 2d). Under control conditions, all lines exhibited a steady increase in leaf number. The WT and *agl16-7* produced consistently more leaves than *agl16-3* and *agl16-10*, which started with a lower leaf number. By SD III, the WT and *agl16-7* had developed 23 and 26 fully expanded leaves, respectively, while *agl16-3* and *agl16-10* had 21 and 22 leaves, respectively. Notably, *agl16-7* displayed a high rate of leaf formation despite its smaller stature. In contrast, stress conditions caused pronounced differences. On SD II, the WT had lost all vital leaves, dropping from the initial count of 17 to zero vital leaves. Consequently, no statistical analyses could be performed on the morphological leaf parameters. In contrast, the knockout lines retained a higher median number of vital leaves, where *agl16-3* retained 3 of 17 vital leaves (18%), *agl16-7* retained 4 of 18 (22%), and *agl16-10* retained 5 of 16 (31%). In the subsequent recovery phase on SD III, all three KO lines retained leaves, with median counts of 5, 4, and 2 fully expanded leaves observed for *agl16-3*, *agl16-7*, and *agl16-10*, respectively. Compared with their respective well-watered controls, they exhibited significantly fewer fully expanded leaves on SD II and SD III (*p* < 0.001). The formation of newly fully expanded leaves was completely inhibited in all WT individuals at both time points. In contrast, the KO lines did not differ significantly from their well-watered controls, with the exception of *agl16-7* at SD II (*p* < 0.05).

Distinct differences were also observed in the analysis of the molecular stress markers MDA and proline. In the WT, the following molecular physiological and genetic leaf analyses were conducted on the few remaining semi-vital leaves (defined as leaves with approx. 60% remaining chlorophyll-filled cells). To gain insight into the experienced oxidative stress, MDA levels were assessed (Fig. 3a). Under control conditions, all four lines exhibited only minor variation in MDA levels among individual plants over time, and remaining below an average of 7.8 nmol g⁻¹ FW. Under stress conditions, all lines displayed elevated MDA levels on SD II, with the stressed WT reaching the highest mean of 15.1 nmol g⁻¹ FW. The WT, *agl16-3* and *agl16-7* showed significant deviations in MDA levels compared to their respective control (*p* < 0.001, *p* < 0.05, *p* < 0.001, respectively), whereas *agl16-10* did not show such deviation. Although the mean MDA was lower in all KO lines than in WT, a significant difference was only observed between the stressed WT and *agl16-10* (*p* < 0.05). On SD III the MDA levels of lines *agl16–3* and *agl16–7*, that had been elevated at SD II, dropped significantly (*agl16–3*: *p* < 0.05; *agl16–7*: *p* < 0.01) to levels comparable with the respective controls. Line *agl16–10*, which did not show an earlier increase, therefore showed no significant change at this SD. In contrast, the WT showed no significant reduction in MDA levels; instead, levels increased further and reached a mean peak of 18.8 nmol g⁻¹ FW, differing significantly from its own control (*p* < 0.001) and from all three KO lines (*p* < 0.001).

**Fig. 3.**
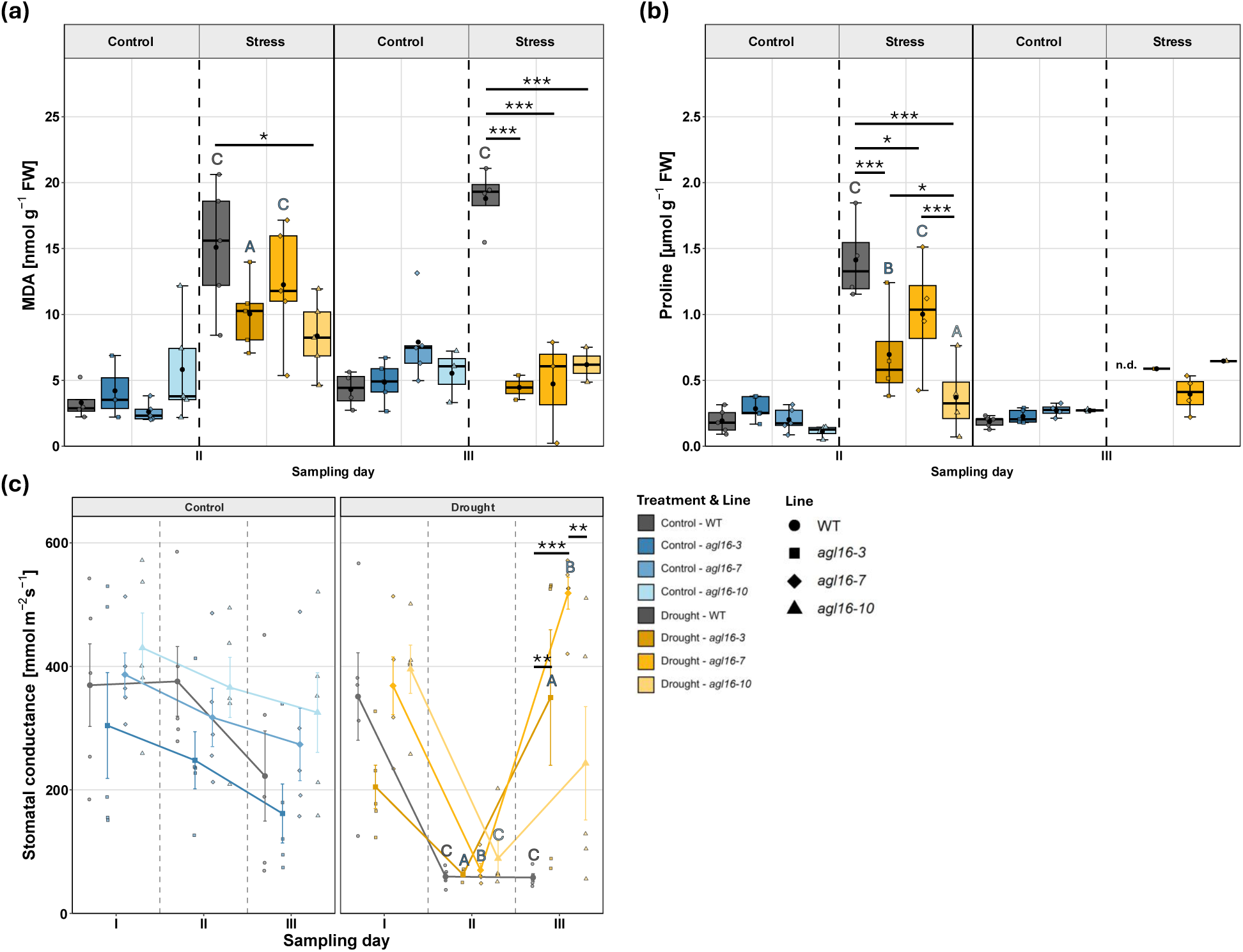
Physiological responses of WT and *PtaAGL16*_2 KO lines (*agl16*-*3, agl16*-*7*, *agl16*-*10*) to drought stress. Measurements of (a) MDA, (b) proline, and (c) stomatal conductance across SDs under control and drought-stress treatments. Data points visualize individual plants (up to *n* = 5 biological replicates per line × treatment × SD). For proline and MDA (A, B), boxplots indicate the median and interquartile range (whiskers: 1.5× IQR), and black dots indicate means. n.d. (not determined) denotes missing proline data, as leaf samples were unavailable. For stomatal conductance (c), lines connect means ± standard error. Asterisks and connecting brackets denote significant pairwise differences among lines within each treatment × sampling day combination based on mixed-effects models followed by Holm-adjusted post hoc contrasts (* *p* < 0.05, ** *p* < 0.01, *** *p* < 0.001). Different capital letters above the stress groups indicate significant treatment contrasts within each line (p < 0.05, p < 0.01, p < 0.001, indicated by A, B, C, respectively). Letters are colored according to the respective control group of each line

Under control conditions, all lines maintained proline levels below a threshold of 0.28 µmol g⁻¹ fresh weight (FW) throughout the experiment (Fig. 3b). In contrast, the stress treatment induced significant changes. At SD II, drought-treated lines of WT, *agl16-3*, *agl16-7* and *agl16-10* exhibited a significant increase in proline levels compared to their respective controls, with mean values of 1.4, 0.7, 1, and 0.4 µmol g⁻¹ FW, respectively (*p* < 0.001, *p* < 0.01, *p* < 0.001, and *p* < 0.05). Notably, proline levels of the WT were significantly higher than those in *agl16-3*, *agl16-7,* and *agl16-10* (*p* < 0.001, *p* < 0.05, and *p* < 0.001, respectively). Among the KO lines, *agl16-3* and *agl16-7* exhibited significantly higher proline levels than *agl16-10* (*p* < 0.05, *p* < 0.001). During the recovery phase, proline levels in the three KO lines decreased, consistent with alleviation of stress. WT proline could not be determined at SD III because viable leaf tissue was no longer available after severe defoliation.

Stomatal conductance was measured as a physiological parameter reflecting stomatal aperture and gas exchange (Fig. 3c). At the beginning of the experiment, the WT, *agl16-7*, and *agl16-10* lines exhibited values ranging from 350 to 420 mmol m⁻² s⁻¹ across both treatments. Only line *agl16-3* displayed lower initial values (200–300 mmol m⁻² s⁻¹), although not significantly different from the other lines. Under control conditions, a general decline in stomatal conductance was observed over the course of the experiment. These differences were not significant between lines or between SDs. In contrast, the stressed lines showed a marked reduction in stomatal conductance during severe stress. Under drought, stomatal conductance decreased significantly in all four lines compared with their respective controls (WT: *p* < 0.001; *agl16-3*: *p* < 0.05; *agl16-7*: *p* < 0.01; *agl16-10*: *p* < 0.001). Mean values at SD II were 60, 63, 70, and 88 mmol m⁻² s⁻¹ for WT, *agl16-3*, *agl16-7*, and *agl16-10*, respectively. No significant differences were detected among the four stressed lines. Following re-watering on SD III, all three KO lines displayed a significant increase in stomatal conductance (*agl16-3* and *agl16-7*: *p* < 0.001; *agl16-10*: *p* < 0.05). Lines *agl16-3* and *agl16-7* reached particularly high mean values (350 and 519 mmol m⁻² s⁻¹, respectively), where both differed significantly from their own control group (*p* < 0.05, *p* < 0.01, respectively), whereas *agl16-10* showed a moderate mean increase to 243 mmol m⁻² s⁻¹. In contrast, the WT showed no significant change in stomatal conductance after re-watering. Consequently, *agl16-3*, and *agl16-7* differed significantly from the WT (*p* < 0.01, *p* < 0.001, respectively). Line *agl16-7* also differed from *agl16-10* (*p* < 0.01).

At the start of the experiment, all individuals in the stress group appeared vital, displaying turgid green leaves and vital shoot tips (Fig. 4). After seven days without water (SD II), the phenotype changed drastically. Nearly all individuals exhibited clear stress symptoms, including leaf and shoot tip wilting. These symptoms were most pronounced in the WT, which displayed wilting shoot tips in all individuals, whereas the KO lines generally presented milder visual symptoms. The extent of stress damage became fully apparent after re-watering. The WT exhibited the most pronounced stressed phenotype across all lines, characterized by dead shoot tips with only residual chlorophyll in the lower stem sections, and no vital leaves remaining. In contrast, the KO lines showed varying degrees of resilience. Line *agl16-3* had three completely vital individuals and two with dead shoot tips but substantial stem chlorophyll. Line *agl16-7* included two individuals that showed similar growth to the controls, and three with dead shoot tips that were already resprouting and had retained some leaves. Finally, *agl16-10* displayed two completely vital individuals, one severely stressed individual with a vital shoot tip, and two individuals with dead shoot tips, although a substantial amount of stem chlorophyll.

**Fig. 4.**
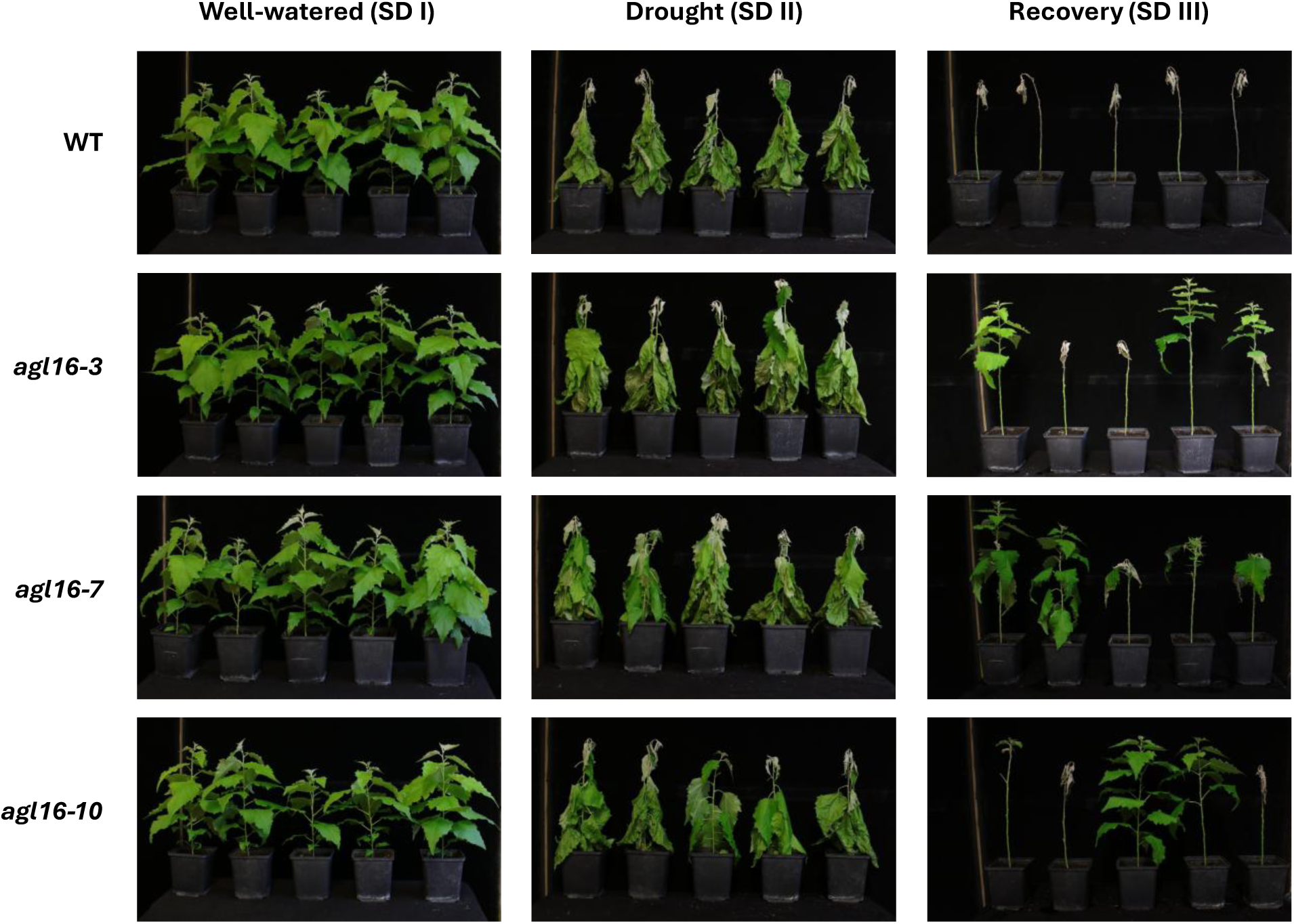
Phenotypic variation of drought-treated poplars in the climate chamber experiment. All individuals (n = 5 per line) of WT and *PtaAGL16_2* KO lines (*agl16-3*, *agl16-7*, and *agl16-10*) are shown at three sampling days: well-watered (SD I), after 7 d without watering (drought, SD II), and after 14 d of re-watering (recovery, SD III). The apparent absence of some leaves is due to prior removal for invasive measurements

These differences in visual stress symptoms are statistically supported. Here, the most pronounced difference between WT and KO lines was observed in shoot vitality after recovery, where all three KO lines retained substantially higher vitality than WT (Fig. 5a). Shoot vitality, calculated as the ratio of vital stem length to total plant height, showed the most severe decline in the WT, with a loss exceeding 65%. In contrast, the KO lines exhibited significantly lower reductions of approx. 14%, 11%, and 18% for *agl16-3*, *agl16-7*, and *agl16-10*, respectively. Consequently, shoot vitality in all three KO lines differed significantly from that of the WT (*p* < 0.001 for all lines). Furthermore, the stressed WT differed significantly from its own control (*p* < 0.001), whereas no such significant differences were observed within the three KO lines.

**Fig. 5.**
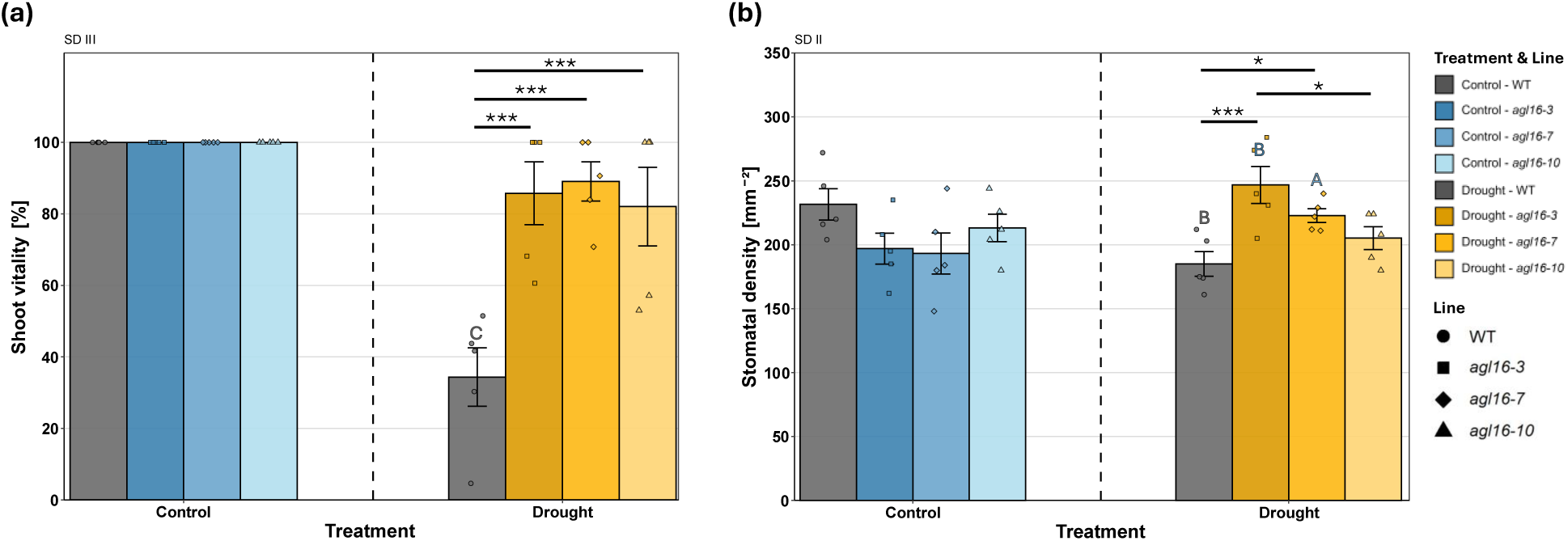
Shoot vitality and stomatal density of WT and *PtaAGL16_2* KO lines (*agl16-3*, *agl16-7*, *agl16-10*) in the drought stress experiment. (a) Shoot vitality assessed at SD III and (b) stomatal density assessed at SD II under control and drought-stress treatments. Bars show means ± standard error; dots indicate individual plants (n = 5 per line × treatment). Asterisks indicate significant pairwise differences among lines within each treatment based on beta regression (vitality) or linear models (stomatal density) followed by Holm-adjusted post hoc contrasts (* *p* < 0.05, *** *p* < 0.001). Different capital letters above the stress groups indicate significant treatment contrasts within each line (p < 0.05, p < 0.01, p < 0.001, indicated by A, B, C, respectively). Letters are colored according to the respective control group of each line

Stomatal density was also determined because AGL16 has been associated with stomatal development in *A. thaliana* (Fig. 5b). Although not statistically significant under control conditions, the WT exhibited a higher mean stomatal density (231 stomata mm⁻²) compared to the KO lines (197, 193, and 213 stomata mm⁻² for *agl16-3*, *agl16-7*, and *agl16-10*, respectively). Here, the comparison between WT and line *agl16-7* showed a marginal non-significant difference (*p* = 0.055). Under severe stress, the WT showed a lower stomatal density of approx. 20%, with 185 stomata mm⁻² (*p* < 0.01). In contrast, *agl16-3* and *agl16-7* displayed a higher stomatal density under stress. Specifically, *agl16-3* showed an increase of 50 stomata per mm⁻² (+25%; *p* < 0.01) and *agl16-7* an increase of 30 stomata per mm⁻² (+15%; *p* < 0.05), compared to their respective controls. Line *agl16-10* exhibited nearly the same stomatal density with slightly less stomata per mm⁻² (–4%). Consequently, significant differences were observed between the lines: *agl16-3* and *agl16-7* differed significantly from the WT (*p* < 0.001 and *p* < 0.05, respectively), and *agl16-3* also differed significantly from *agl16-10* (*p* < 0.05).

To assess the molecular response, we analyzed the expression of several genes via RT-qPCR. Under drought stress, transcript levels of all assayed targets responded largely in a treatment-dependent manner. The transcript abundance of candidate gene *PtaAGL16_2* was increased under drought in the WT and all mutant lines (Fig. 6a). Relative to their own controls, *PtaAGL16_2* abundance was increased in all stress-treated lines (WT: fold change (FC) = 12.2, *agl16-3*: 5.5, *agl16-7*: 8.0, and *agl16-10*: 5.6; with *p* < 0.001 for all lines). Direct comparison under drought further indicated significantly lower abundance in the mutant lines than in the WT (*p* < 0.01 for all lines The unedited paralog *PtaAGL16_5* was also induced by drought (Fig. 6b). All lines exhibited a drought-induced increase compared to their own controls (WT: FC = 53.3 (*p* < 0.001), *agl16-3*: 5.7 (*p* < 0.01), *agl16-7*: 44.6 (*p* < 0.001), and *agl16-10*: 4.0 (*p* < 0.05)). Direct comparison under drought revealed significantly lower expression in the KO lines *agl16-3*, *agl16-7*, and *agl16-10* compared with the WT (*p* < 0.01, *p* < 0.05 and *p* < 0.05, respectively). Under control conditions, the WT, *agl16-3* and *agl16-10* differed significantly from *agl16-7* (*p* < 0.05 for all lines).

**Fig. 6.**
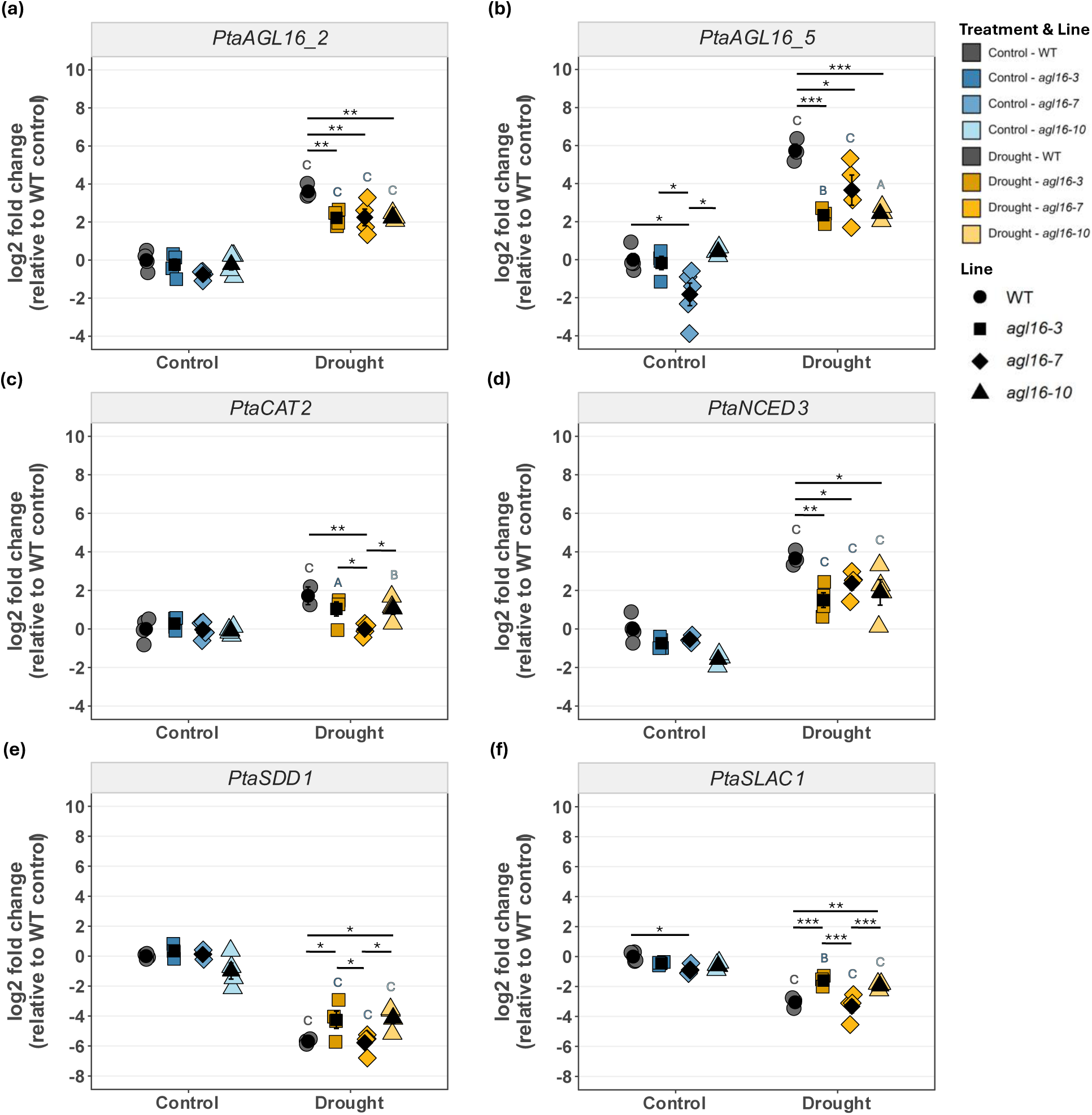
Gene expression analyses of WT and *PtaAGL16_2* KO lines (*agl16-3*, *agl16-7*, and *agl16-10*). Measurements of (a) *PtaAGL16_2*, (b) *PtaAGL16_5*, (c) *PtaCAT2*, (d) *PtaNCED3*, (e) *PtaSDD1*, and (f) *PtaSLAC1* under control and drought-stress treatment. Colored data points visualize individual plants (n = 3-4 biological replicates per line × treatment). Expression is shown as log2 fold change (relative to WT control). Black data points indicate means. Asterisks and connecting lines denote significant differences within each treatment based on linear models followed by FDR-adjusted post hoc contrasts (* *p* < 0.05, ** *p* < 0.01, *** *p* < 0.001). Different capital letters above the stress groups indicate significant treatment contrasts within each line (p < 0.05, p < 0.01, p < 0.001, indicated by A, B, C, respectively). Letters are colored according to the respective control group of each line

The antioxidant defense-related gene *PtaCAT2* was significantly induced in all stressed-treated lines except for line *agl16-7* (WT: FC = 3.3 (*p* < 0.001), *agl16-3*: 1.69 (*p* < 0.05), *agl16-7*: 1.0, and *agl16-10*: 2.2 (*p* < 0.01)) (Fig. 6c). Comparison under drought revealed a significantly lower expression in the KO *agl16-7* compared to WT, as well as to *agl16-3*, and *agl16-10* (*p* < 0.01, *p* < 0.05, and *p* < 0.05 respectively).

We additionally analyzed the expression of *PtaNCED3*, which takes part in the ABA biosynthesis (Fig. 6d). All lines significantly increased the *PtaNCED3* gene expression upon the drought treatment in comparison to their respective controls (WT: FC = 12.7, *agl16-3*: 4.7, *agl16-7*: 7.5, and *agl16-10*: 11.2; *p* < 0.001 for all lines). Between-line comparisons under the control treatment revealed no significant expression differences. Under the drought treatment, we found a significant lower *NCED3* expression *of agl16-3*, *agl16-7*, and *agl16-10* compared to the WT (*p* < 0.01, *p* < 0.05, and *p* < 0.05, respectively).

We also analyzed *PtaSDD1* expression, as *SDD1* has been associated with stomatal density and distribution in *A. thaliana* (Fig. 6e). Upon drought stress, all lines shared the significant downregulation of *PtaSDD1* in comparison to their respective controls (WT: FC = 0.02, *agl16-3*: 0.04, *agl16-7*: 0.02, and *agl16-10*: 0.12; *p* < 0.001 for all lines). Further, direct comparisons under the drought treatment revealed a significant lower expression of the WT and *agl16-7* compared to *agl16-3* and *agl16-10* (*p* < 0.05 for all comparisons).

*PtaSLAC1*, which is related to ABA-induced stomatal regulation, showed a significantly reduced expression under drought treatment in all lines compared to their respective controls (WT: FC = 0.12 (*p* < 0.001), *agl16-3*: 0.45 (*p* < 0.01), *agl16-7*: 0.18 (*p* < 0.001), and *agl16-10*: 0.39 (*p* < 0.001)) (Fig. 6f). Direct comparison between lines under control treatment revealed a significantly lower gene expression in *agl16-7* compared to WT (*p* < 0.05). Under the drought treatment, *agl16-3* showed a significantly higher expression compared to WT (*p* < 0.001), and *agl16-7* (*p* < 0.001). Further, *agl16-10* also showed a significantly higher expression than WT (*p* < 0.01) and *agl16-7* (*p* < 0.001).

## Discussion

In the context of improving plant stress resilience, TFs represent key regulatory components by integrating environmental signals and modulating downstream gene expression networks, especially in the control of complex traits such as drought tolerance. Generally, the MADS-box TF family is well-characterized in plants, with members involved in a wide spectrum of processes, including plant growth, reproductive development, and abiotic and biotic stress responses (Zhang et al. 2024). While the genomic landscape of this family has also been mapped in woody species like poplar (Leseberg et al. 2006), the validation of gene functions remains scarce. Specifically, recent work in model systems has elucidated the function of the MADS-box TF AGL16 in diverse processes, including stomatal development, flowering time, and abiotic stress responses to drought and salinity (Zhao et al. 2020; Zhao et al. 2021; Dong et al. 2023). Although AGL16 has also been investigated in perennial cotton, its role in long-lived woody tree species, which generally face distinct physiological constraints due to their long lifespan and large size, remains unknown. To address this, we investigated the role of PtaAGL16 in hybrid poplar using a CRISPR/Cas9 KO approach to assess its contribution to drought resilience, including both stress tolerance and post-stress recovery capacity.

### Morphological and physiological drought responses across tested lines

Plants exposed to drought rely on a combination of avoidance, tolerance, and recovery mechanisms (Berger et al. 2016). In response to increasing water deficit, plants sequentially activate physiological strategies to maintain function or ensure survival. Stomatal closure is a key mechanism of dehydration avoidance, helping to reduce water loss during moderate drought and as stress intensifies, growth ceases and other strategies such as cavitation tolerance become critical for long-term survival (Volaire 2018). Among these strategies, there is also dehydration escape, which comprises completing the life cycle before the onset of severe water shortage, that is predominantly observed in annuals (Franks 2011; Kooyers 2015). In contrast, perennial woody species like poplars cannot employ this escape strategy and must rely on drought avoidance/tolerance and recovery mechanisms to survive recurring drought periods. From a morphological standpoint, drought avoidance can be enhanced by traits such as reduced height and smaller total leaf area (achieved via fewer or smaller leaves). This is because such traits reduce transpirational water loss while maintaining essential physiological processes (Toscano et al. 2019). Consistent with these endurance mechanisms, our data show growth arrest across all stress-treated lines during peak stress at SD II. Although the full KO lines *agl16-7* and *agl16-10* generally tended to exhibit reduced growth, these morphological differences alone are unlikely to account for their enhanced drought resilience. In particular, *agl16-3* closely resembled WT in plant height, whereas *agl16-7* produced the highest number of fully expanded leaves, although this difference was not statistically significant. Thus, the absence of a consistent relationship between constitutive growth traits and drought resilience suggests that the enhanced resilience of the KO lines is more likely associated with altered stress-responsive regulation. In parallel, a strong reduction of stomatal conductance from approx. 330 to 70 mmol m⁻² s⁻¹ could be observed in all stress-treated lines and is consistent with a strong stress response in poplar (Yu et al. 2021). Interestingly, control plants also displayed a gradual decline in stomatal conductance over the course of the experiment. Because the control plants were well watered and relative humidity was controlled, soil water deficit is unlikely to explain this decline. Although the underlying mechanism remains unclear, one possible explanation is plant-to-plant signaling mediated by airborne cues. Previous studies have shown that volatile organic compounds and other airborne signals can modulate stomatal behavior in neighboring plants exposed to stress (Caparrotta et al. 2018; Jin et al. 2021).

### Preserved shoot vitality supports the superior post–drought recovery in *agl16* KO lines

Beyond these physiological responses, the most pronounced difference between the *agl16* mutants and WT became apparent during recovery after re-watering. The enhanced drought resilience of the mutants was most evident in their superior recovery capacity, characterized by the resumption of growth parameters and the formation of new leaves and leaf buds following re-watering. A prerequisite for shoot recovery is the preservation of viable meristematic tissues, particularly at the shoot apex. As the primary site of shoot organogenesis and a central regulator of shoot development, the shoot apical meristem represents a key tissue for maintaining growth potential under changing water availability (Lafon-Placette et al. 2018). Preserving the viability of this tissue may therefore facilitate the rapid resumption of shoot growth following re-watering.

In our experiment, the WT suffered complete defoliation and severe stem necrosis, whereas the *agl16* mutants retained substantial foliage. The initial leaf shedding from the lower stem observed in all lines may reflect a resource reallocation strategy, consistent with reports from young poplars that mobilize nutrients from older leaves to sustain developing tissues (Yao et al. 2024). Whereas defoliation progressed to complete leaf loss in the WT, it remained incomplete in the KO lines. Thus, retention of upper foliage may have helped maintain photosynthetic capacity during recovery, potentially facilitating the resumption of shoot growth after re-watering.

Furthermore, shoot vitality, defined here as the percentage of chlorophyll-containing stem tissue, was significantly higher in the *agl16* mutants (approx. 86%) compared to the WT (34%), reaching levels comparable to well-watered controls. The persistent stem necrosis and lack of regrowth in the WT indicate more severe drought-induced shoot damage. Such damage may have constrained the restoration of shoot function, potentially including hydraulic recovery, and consequently limited the resumption of shoot growth after re-watering (Zwieniecki and Secchi 2015; Secchi et al. 2017). In contrast, the retention of chlorophyll-containing stem tissue in the KOs may provide a greater potential for stem-associated photosynthetic carbon assimilation, including cuticular photosynthesis, which has been proposed to support local carbon balance and vascular maintenance during drought (Bloemen et al. 2016; Baerdemaeker et al. 2017; Ávila-Lovera et al. 2024). Whether this contributes to the observed recovery advantage remains to be determined.

### Reduced cellular stress and attenuated stress-responsive gene activation in *agl16* mutants

The observed divergence in recovery raises the possibility that, despite similar acute stress responses, the *agl16* mutants experienced a lower cellular stress burden than the WT. Under drought, plants face coupled osmotic and oxidative stress, leading to oxidative damage of cellular membranes. Here, MDA is commonly used as an indicator of lipid peroxidation and associated oxidative damage (Del Rio et al. 2005; Morales and Munné-Bosch 2019). Although MDA increased in all drought-treated lines, levels remained elevated exclusively in the WT under re-watering conditions. In strong contrast, MDA levels in all *agl16* mutants returned to levels comparable to those of well-watered controls, suggesting that the elevated lipid peroxidation observed during drought was largely resolved during recovery. The persistence of elevated MDA levels in the WT after re-watering suggests that oxidative damage was not resolved during recovery.

In addition to oxidative damage, drought responses can also be assessed through changes in compatible solutes such as proline. In annual crops, higher proline accumulation has been shown to increase drought stress tolerance (Furlan et al. 2020; Shrestha et al. 2022). In contrast, in woody perennials including *Populus* spp., proline accumulation has more frequently been associated with drought severity than with drought tolerance. Thus, proline is here best interpreted primarily as a stress marker rather than a determinant of drought tolerance (Rennenberg et al. 2006; Shvaleva et al. 2006; Fan et al. 2025). This interpretation is further supported by our results, as the WT exhibited significantly higher proline contents than all *agl16* mutants while simultaneously displaying the most severe drought-stressed phenotype. Together, the elevated MDA and proline levels in the WT indicate a significantly higher internal stress burden compared to the *agl16* mutants, which is further supported by the gene expression analysis of *PtaCAT2* and *PtaNCED3*.

A common plant response to drought involves the upregulation of the antioxidant defense system to detoxify reactive oxygen species (ROS), as well as the accumulation of the phytohormone ABA to minimize water loss through stomatal closure (Haghpanah et al. 2024). *PtaCAT2*, encoding a peroxisomal catalase that decomposes H₂O₂ generated largely during photorespiration, was more strongly induced in the WT, consistent with increased ROS pressure and/or activation of compensatory antioxidant defenses under water deficit (Du et al. 2008). *PtaNCED3*, encoding the rate-limiting enzyme in stress-induced ABA biosynthesis, was likewise more strongly upregulated in the WT, suggesting stronger activation of ABA biosynthesis under drought stress (Iuchi et al. 2001; Lee et al. 2021; Yu et al. 2021). Together, these indicators point to a stronger and/or more persistent cellular stress response in the WT compared with the mutants, which showed preserved shoot vitality and superior recovery.

### Changes in stomatal traits and expression of ABA/stomatal-associated genes

Based on findings in *A. thaliana*, it could be assumed that AGL16-mediated regulation of *SDD1* expression is correlated to reduced stomatal density (Zhao et al. 2020). Under well-watered conditions, the stomatal density of all three KO lines was 7-16 % lower than in the WT, although this difference was not statistically significant.

Under drought, all lines showed strong downregulation of *PtaSDD1*, yet stomatal density diverged, decreasing in the WT but increasing in *agl16-3* and *agl16-7*. This divergence, together with the modest differences observed under control conditions, suggests that stomatal patterning may be influenced by factors beyond PtaSDD1 expression, including altered leaf expansion and systemic signals from mature leaves (Miyazawa et al. 2006; Xu and Zhou 2008), and does not support a simple relationship between *PtaAGL16* disruption and stomatal density, as also suggested for *GhAGL16* overexpression in *A. thaliana* (Lei et al. 2024b). A direct causal link between reduced stomatal density and the reduced stature observed in *agl16-7* and *agl16-10* cannot be established, as this relationship was not consistent across all lines.

The expression of selected ABA-associated genes (*PtaNCED3*, *PtaSLAC1*) differed quantitatively between WT and *agl16* lines under drought. In *A. thaliana*, *agl16* mutants exhibit accelerated stomatal closure under drought, accompanied by transcriptional changes in ABA-biosynthesis genes, suggesting that AGL16 acts as a negative regulator of ABA-mediated drought responses (Zhao et al. 2020). Similarly, silencing of *GhAGL16* in *G*. *hirsutum* enhanced drought tolerance and was associated with ABA-responsive transcriptional changes (Lei et al. 2024a). Although ABA contents were not measured in this study, the lower *PtaNCED3* expression in all KO lines indicates altered transcriptional regulation of ABA biosynthesis. Under drought stress, ABA signalling results in the phosphorylation of SLAC1, thereby promoting anion efflux from guard cells, and ultimately rapid stomatal closure (Geiger et al. 2009). While *SLAC1* transcription expression has been reported to increase during the early phase of drought stress, supporting the establishment of the stomatal closure response, prolonged drought is frequently associated with reduced *SLAC1* transcript abundance (Mo et al. 2023). Under prolonged drought, sustained stomatal closure relies primarily on ABA-dependent activation and post-translational regulation of pre-existing guard-cell signalling components, including activation of SLAC1, rather than requiring continued transcriptional induction of SLAC1 (Geiger et al. 2009). In our study, *SLAC1* was downregulated in all lines under drought, although the magnitude differed between the lines. Whether these transcriptional changes reflect altered ABA levels or ABA sensitivity, and whether PtaAGL16_2 directly regulates ABA- and stomatal-associated genes, remains to be experimentally validated.

The observation that *PtaAGL16_2* transcript levels are induced by drought in WT plants, yet its disruption enhances drought resilience, suggests that PtaAGL16_2 may negatively influence drought adaptation in hybrid poplar. The functional significance of drought-induced *PtaAGL16_2* expression and the mechanism by which its loss improves drought resilience remain to be studied.

### Limitations and future directions

Despite the presence of the *PtaAGL16* paralog on chromosome 5, the disruption of the chromosome 2 copy alone was associated with a pronounced drought-resilient phenotype. Notably, line *agl16-3*, which carries an in-frame deletion on one allele, resulting in the loss of a single amino acid, displayed an intermediate phenotype between the other KO lines and the WT but still exhibited markedly improved drought recovery. The consistent direction of the drought-recovery phenotype across three lines with distinct indels supports the specific role of

PtaAGL16_2, although the magnitude of the phenotype did not show a simple relationship with the predicted molecular consequences of the individual indels. No evidence for the assessed off-target events was detected, supporting the interpretation that the observed phenotypes are associated with the intended gene modification. The consistent phenotype across three independently edited lines with distinct indels and independent T-DNA integration sites strongly supports the conclusion that the observed drought-recovery phenotype is attributable to the disruption of *PtaAGL16_2*.

Together, these findings support a role of *PtaAGL16_2* in limiting drought recovery in hybrid poplar, although functional redundancy with the chromosome 5 paralog cannot be excluded. The chromosome 5 paralog, *PtaAGL16_5*, was not functionally investigated because the available gRNA resulted in insufficient editing efficiency. Functional disruption of *PtaAGL16_5*, either alone or in combination with *PtaAGL16_2*, would therefore provide a useful approach to determine whether the two paralogs have redundant functions and whether complete loss of AGL16 activity further alters drought performance or affects plant growth under non-stress conditions.

Although the proposed regulatory network is supported by transcriptional and physiological evidence, these data do not establish a direct regulatory mechanism. ABA quantification and the direct involvement of *PtaAGL16_2* in the regulation of ABA- and stomatal-associated genes by *PtaAGL16_2* remains to be experimentally validated. Future work should address whether the observed drought resilience is stable across environments and growing seasons and whether it is associated with trade-offs in growth, biomass production, or responses to additional abiotic stresses.

## Conclusion

Overall, CRISPR/Cas9-mediated disruption of *PtaAGL16_2* is associated with enhanced drought recovery in hybrid poplar, accompanied by multiple physiological and molecular changes. The improved recovery of the edited lines was associated with reduced oxidative stress indicators, preserved shoot vitality, and altered expression of ABA- and stomatal-associated genes. Together, these responses are consistent with improved maintenance of tissue viability during drought and enhanced recovery capacity after re-watering. Collectively, these findings highlight *PtaAGL16* as a potential target for genome editing and molecular breeding of climate-resilient forest trees, particularly in breeding programs targeting drought resilience in fast-growing woody crops.

## Supporting information

Online Resource

## Acknowledgements

We are grateful to Susanne Jelkmann and Ian Volkmann for excellent technical support.

## Data Availability Statement

The original contributions presented in this study are included in the article and Supplementary Materials. Further inquiries can be directed to the corresponding author.

## Author Contributions

AF, MF, and TB designed the research. AF and SW performed the research and analyzed the data. AF and TB interpreted the data. AF and TB wrote the manuscript, and all the authors edited the text. MF and TB received the funding.

