## Supplementary material for "CRISPR/Cas9 knockout of *PtaAGL16* improves post-drought recovery and shoot vitality in hybrid poplar": Online Resource

Fendel A, Wodtke S, Fladung M, Bruegmann T*
*) Thünen Institute of Forest Genetics, Grosshansdorf, Germany;

#### Supplemental Material

**(a)**

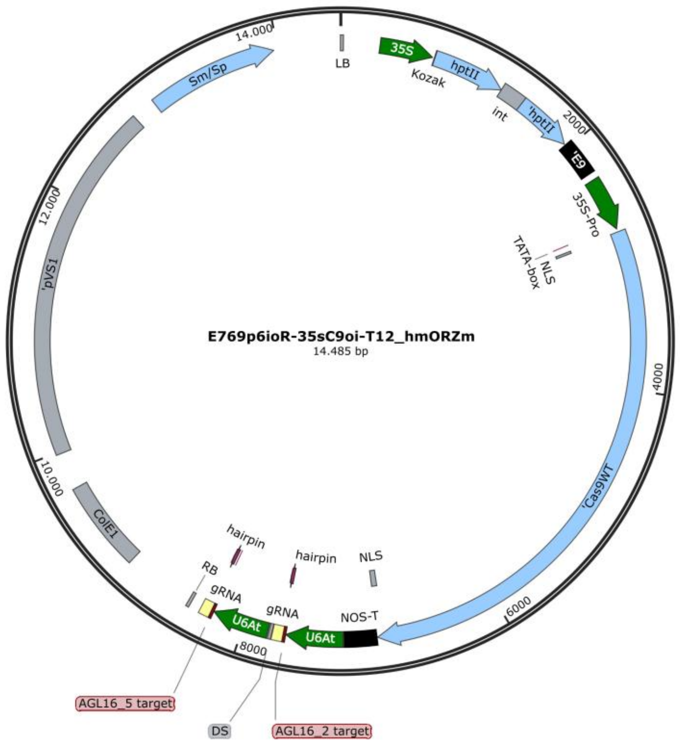


**(b)**

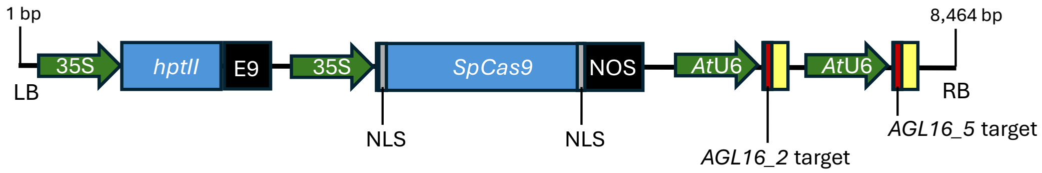


**Fig. S1:** Transformation vector used for generation of *agl16* mutant lines in *Populus tremula* × *P. alba*. (a) Illustration of the total vector including the T-DNA and vector backbone, with a total size of 14,485 bp. The vector backbone consists of a *Colicin E1 Origin of Replication* (*ColE1* origin of replication), a *Plasmid pVS1 Origin of Replication*, and a *Streptomycin/Spectinomycin* resistance gene. (b) From the left border (LB) to the right border (RB) of the T-DNA, the vector contains a *Cauliflower Mosaic Virus 35 Promoter* (*CaMV 35S* promoter), a *HYGROMYCIN PHOSPHOTRANSFERASE II* gene (*hptII* resistance gene), a terminator from *Pisum sativum* (*E9* terminator), another *CaMV 35S* promoter, driving the expression of the *SpCas9* gene fused to both an N- and a C-terminal nuclear localization signal (NLS), followed by a *NOPALINE SYNTHASE* terminator (*NOS* terminator), and the two gRNAs containing the respective *AGL16* spacers, driven by the *A. thaliana* U6 small nuclear RNA (U6) promoter.

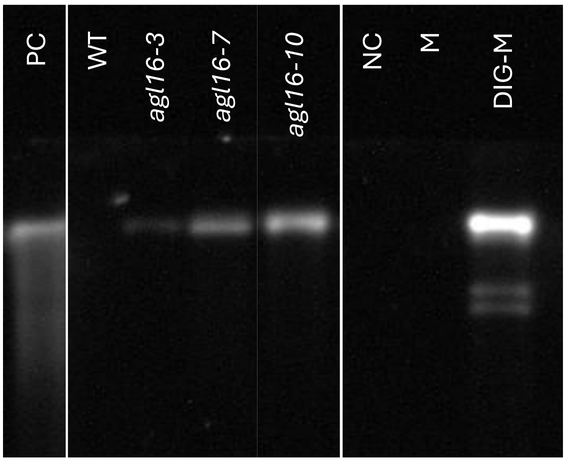


**Fig. S2:** Southern blot analysis of T-DNA insertion frequencies in transgenic lines. Genomic DNA was digested with *Dra*I, separated by agarose gel electrophoresis and transferred to a nylon membrane. The membrane was hybridized with a digoxigenin (DIG)-labeled probe specific to the *HPTII* gene. Lane 1: Positive control (plasmid containing the *HPTII* sequence); Lane 2: WT; Lanes 3-5: *agl16-3*, *agl16-7*, *agl16-10*; Lane 6: Negative control ($H_2O$); Lane 7: DNA molecular weight marker; Lane 8: DIG-labeled DNA marker

#### *PtaAGL16_2* (PtXaTreH.02G093900, CDS WT, *P. tremula* allele)

5'-ATGGGGAGGGGGAAGATAGTGATAAGAAGGATTGACAACTCGACAAGCAGGCAAGTGACCTTCTCAAAGAGG
AGGAATGGATTGTTGAAGAAAGCGAAAGAGCTGGCGATCCTTTGCGACGCAGAAGTTGGAGTCATGATCTTCTCT
AGCACCGGCAAGCTCCATGATTTCTCCAGCACCAGCATGAAATCAGTCATTGAAAGATACAACAAATCAAAAGAC
GAGCATCATCAAATGGGCAACCCAACCTCTGAAGTCAAGTTCTGGCAAAGGGAGGCAGCAGTGTTGAGACAGCAA
CTGCAAACCTTGCAAGAAAATCATCGGCAAATGATGGGCGAACAACTTTCTGGCTTAAGCATAACAGATTTGCAG
AATTTGGAGAACCAATTGGAAATGAGTCTTCAGGGTGTTCGCATGAAAAAGGACCAAATTTTAATGGATCAAATA
CAAGAACTAAATAGAAAGGGCAACCTCATTCACCAAGAAAATGTTGAACTCTATCAGAAGGTCTATGGAACAAGG
GACGTAAACAGAGCAAACAGAATTTCCCTTCTCACAAACGGTCTGGCCATCGGAGAGGAATCACATGTACCTGTC
CATCTCCAGCTTAGCCAGCCACAACAACAAAACCATGATACACCAGCTACAAAGTTGGGGTACAATTTCCTCCAC
ATCACAAGTATTTAA-3'

#### *PtaAGL16_2* (PtXaAlbH.02G094700, CDS WT, *P. alba* allele)

5'-ATGGGGAGGGGGAAGATAGTGATAAGAAGGATTGACAACTCGACAAGCAGGCAAGTGACCTTCTCAAAGAGG
AGGAATGGATTGTTGAAGAAAGCGAAAGAGCTGGCGATCCTTTGCGACGCAGAAGTTGGAGTCATGATCTTCTCT
AGCACCGGCAAGCTCCATGATTTCTCCAGCTCCAGCATGAAATCAGTCATTGAAAGATACAACAAATCAAAAGAC
GAGCATCATCAAATGGGCAACCCAACCTCTGAAGTCAAGTTCTGGCAAAGGGAGGCAGCAGTGTTGAGACAGCAA
CTGCAAACCTTGCAAGAAAATCATCGGCAAATGATGGGCGAACAACTTTCTGGCTTAAGCATAACAGATTTGCAG
AATTTGGAGAACCAATTGGAAATGAGTCTTCAGGGTGTTCGCATGAAAAAGGACCAAATTTTAATGGATCAAATA
CAAGAACTAAATAGAAAGGGCAACCTCATTCACCAAGAAAATGTTGAACTCTATCAGAAGGTCTATGGAACAAGG
GACGTAAACAGAGCAAACAGAATTTCCCTTCTCACAAATGGTCTGGCCATCGGAGAGGAATCACATGTACCTGTC
CATCTCCAGCTTAGCCAGCCACAACAACAAAACCATGATACACCAGCTACAAAGTTGGGGTACAATTTCCTCCAC
ATCACAAGTATTTAA-3'

**Fig. S3**: Coding sequences (CDS) of the candidate gene *PtaAGL16*_2 from both WT alleles. The gRNA spacer sequences used for CRISPR/Cas9 targeting are highlighted in green, and the corresponding PAM sequences are shown in red

### Supplemental Table S1

### Primers used for molecular analyses

| Forward (5' - 3') | Reverse (5' - 3') | Primer binding site(s) | Gene identifier in *P. tremula* × *P. alba* | Application |
|---|---|---|---|---|
| GAGAAGTTTGATAGCGTGTCTG | TAGCGTCACAGCGGCCTTG | *hptII* | | genotyping of transformed antibiotic resistance |
| GCTCCAGACAAGAAGTACAGC | TGTTCACGCGAAGGATGTCG | *SpCas9* | | genotyping of transformed *SpCas9* gene |
| ATATAGCCTCTTTGCTGATGTGC | CTGGAGAAATCATGGAGCTTGC | *PtaAGL16_2* | PtXaTreH.02G093900<br>PtXaAlbH.02G094700 | Target PCR for Sanger sequencing |
| TGGATCACCATCTCTTCCTTG | CACAACCATAAGAAGCTAGCTTG | *PtaAGL16_5* | PtXaTreH.05G115600<br>PtXaAlbH.05G117500 | Target PCR for Sanger sequencing |
| WGTGNAGWANCANAGA | WGTGNAGWANCANAGA | randomly in the genome, 256-fold degenerated | | TAIL-PCR |
| / | CGTGAAGTTTCTCATCTAAGCC | T-DNA in the direction of LB | | TAIL-PCR |
| / | TCGCCTATAAATACGACGGATCG | T-DNA in the direction of LB | | TAIL-PCR |
| / | AGATTTCCCGGACATGAAGCC | T-DNA in the direction of LB | | TAIL-PCR |
| CCAGCCGAGAAGGTCTCCATC | GAACTCGTCAAGCAGGCGGTAG | *nptIIo* | | Southern Blot probe |
| GTCCACCCTCCATTTGGTCC | GGTCTTTCCTGTGAGCGTCT | *PtaUBQ3/4* | PtXaTreH.01G338500<br>PtXaAlbH.01G340500 | RT-qPCR |
| AGGAATGGATTGTTGAAGAAAGCG | CTGGAGAAATCATGGAGCTTGC | *PtaAGL16_5* | see above | RT-qPCR |
| TTCTCAAAGAGGAGGAGTGGG | CATGCTGGTGCTAGAGAAATCATA | *PtaAGL16_5* | see above | RT-qPCR |
| TGCTCCCAAGTGTGCTCATC | TGACCTCCTCATCCCTGTGC | *PtaCAT2-like* | PtXaTreH.02G007700<br>PtXaTreH.05G197800<br>PtXaAlbH.02G008100<br>PtXaAlbH.05G199400 | RT-qPCR |
| GACTACTGCAGATGTAACTGACC | CAATGGCGAAAACTCCTGCC | *PtaSDD1* | PtXaTreH.02G211800<br>PtXaAlbH.02G211300 | RT-qPCR |
| GGGTCTCCGGTGATTTATGACAAG | GCAGTCAGGTGTTTCAATCCACC | *PtaNCED3* | PtXaTreH.11G090100<br>PtXaTreH.01G320100<br>PtXaAlbH.11G081500<br>PtXaAlbH.01G321700 | RT-qPCR |
| GTACCCCCTCTGCTAGCAC | GCCATTGGCCATAGATCTTGAGC | *PtaSLAC1* | PtXaTreH.01G095900<br>PtXaAlbH.01G097800 | RT-qPCR |

Fendel A, Wodtke S, Fladung M, Bruegmann T*
*) Thünen Institute of Forest Genetics, Grosshansdorf, Germany;

### Supplemental Table S2

### T-DNA integration sites

| Transgenic line | Haplotype | Chr. | T-DNA insertion site (bp) | Genomic context | Distance to closest gene (bp) | Closest gene (locus ID) | Gene annotation of closest gene |
|---|---|---|---|---|---|---|---|
| ***agl16-3*** | HAP1, *P. tremula* allele | 02 | 16340423 | Intergenic | 8567 | PtXaTreH.02G172600 | E3 UBIQUITIN-PROTEIN LIGASE BAH1 |
| ***agl16-7*** | HAP1, *P. tremula* allele | 05 | 75500 | Intergenic | 607 | PtXaTreH.05G001100 | solute carrier family 15 (peptide/histidine transporter), member 3/4 (SLC15A3_4, PHT) |
| ***agl16-10*** | HAP1, *P. tremula* allele | 17 | 6519628 | Intergenic | 6898 | PtXaTreH.17G047500 | F13K23.8 PROTEIN |
